# Presence of multiple parasitoids decreases host survival under warming, but parasitoid performance also decreases

**DOI:** 10.1101/2021.08.24.457463

**Authors:** Mélanie Thierry, Nicholas A. Pardikes, Benjamin Rosenbaum, Miguel G. Ximénez-Embún, Jan Hrček

## Abstract

Current global changes are reshaping ecological communities and modifying environmental conditions. We need to recognize the combined impact of these biotic and abiotic factors on species interactions, community dynamics, and ecosystem functioning. Specifically, the strength of predator-prey interactions often depends on the presence of other natural enemies: it weakens with competition and interference or strengthens with facilitation. Such effects of multiple predators on prey are likely to be affected by changes in the abiotic environment, altering top-down control, a key structuring force in natural and agricultural ecosystems. Here, we investigated how warming alters the effects of multiple predators on prey suppression using a dynamic model coupled with empirical laboratory experiments with *Drosophila*-parasitoid communities. While multiple parasitoids enhanced top-down control under warming, parasitoid performance generally declined when another parasitoid was present due to competitive interactions. This could reduce top-down control over multiple generations. Our study highlights the importance of accounting for interactive effects between abiotic and biotic factors to better predict community dynamics in a rapidly changing world and thus better preserve ecosystem functioning and services such as biological control.

## Introduction

Ongoing global anthropogenic changes alter the abiotic context, changing the outcome of species interactions [1,2]. Global warming can modify the strength of trophic interactions due to changes in metabolic rates [3], shifts in spatial distributions and seasonal phenology [4], lethal effects on predators, or altered attack rates [5–7]. But warming also alters the strength of non-trophic interactions among predators [8,9]. Altered non-trophic interactions among predators would change the effects of multiple predators on top-down control [10,11], yet to what extent is unclear. Effects of warming on non-trophic interactions among predators are often overlooked but essential to accurately forecast ecological consequences of warming for biological control and ecosystem integrity.

The effects of multiple predators on prey suppression are often not additive. Additivity would occur if predators had independent effects on prey, in which case increased predator density should enhance top-down control because of higher predatory pressure on the prey. However, direct and indirect interactions among predators may cause effects to deviate from additivity [12–14]. The effects of multiple predators on prey can be synergistic (i.e., the effects are greater than what would be expected if they were additive) due to niche complementarity or facilitation (i.e., risk enhancement for the prey) [15]. By contrast, the effects of multiple predators on prey can be antagonistic due to intraguild predation, competition, or interference when the degree of overlap between predator’s foraging areas or phenologies is too high (i.e., risk reduction) [16]. All such potential effects are called multiple predator effects (MPEs [17]). Emergent MPEs are particularly important in biological control where introducing one or several predator species might result in risk reduction for the prey because of competition among predators instead of planned risk enhancement [18].

Warming can alter both trophic and non-trophic interactions. Changes in the strength of these interactions could modify emergent MPEs, either enhancing or decreasing top-down control. Climate change also disrupts species composition of communities [4,19], which would change the outcome of pairwise interactions that are influenced by other species in the community [20–22]. Changes in species composition of communities are thus also likely to alter MPEs, affecting biological control. However, interactive effects between warming and community composition on top-down control remain poorly studied, and little is known about how warming alters the effects of multiple predators on top-down control.

Here, we used mathematical models in combination with a series of three laboratory experiments on *Drosophila simulans* and three of its co-occurring larval parasitoids to investigate the effects of warming on multiple predator effects on top-down control. Host-parasitoid interactions are a particular type of predator-prey interaction in which parasitoid larvae feed and develop inside or on an arthropod host while adults are free-living [23]. When host is parasitized, three outcomes are possible: the parasitoid successfully develops and kills the host, the host survives and successfully eliminates its parasitoid through immune response (i.e., encapsulation and melanization) [24], or both parties die. The presence of multiple parasitoids can result in extrinsic competition between adults for space and oviposition (i.e., interference) and intrinsic competition within a host [25]. Intrinsic competition results from superparasitism or multiparasitism events when two parasitoids (conspecific or heterospecific) parasitize the same host individual. A single parasitoid can also lay multiple eggs in a single host as part of a strategy to overwhelm the host immune system. In solitary parasitoids, such as the species used in the present study, only one individual completes development in each host, suppressing the other(s) physically or physiologically. Parasitoids represent an excellent system to study how warming directly changes the effects of multiple predators on top-down control because the outcome of the interactions is easily observed by rearing the host, and intrinsic competitive interactions between parasitoids can be observed by dissecting the host larva. In this study, we empirically measured trophic interaction strength across temperatures and parasitoid assemblages. We recorded emergent effects of multiple parasitoids on host suppression by comparing empirical data with estimates in which multiple parasitoids would not interact (i.e., would have an additive effect) using a mathematical model for multiple co-occurring parasitoids with a functional response approach [26,27]. With this framework, we addressed three specific questions: (1) Do multiple parasitoids have additive, synergistic, or antagonistic effects on host suppression? (2) To what extent does temperature modify the outcomes of MPEs? (3) Are changes in host immune response or competitive interaction strength causing emergent MPEs? Our results demonstrate the principal role of temperature for non-trophic interactions among parasitoids, with cascading effects on host suppression.

## Materials and Methods

### Biological system

Cultures of *Drosophila simulans* and their associated parasitoids collected from two tropical rainforest locations in North Queensland Australia: Paluma (S18° 59.031’ E146° 14.096’) and Kirrama Range (S18° 12.134’ E145° 53.102’; both <100 m above sea level; [28]) were used for the experiments. Tropical species already live close to their upper thermal limits [29]. *Drosophila* species are limited in their evolutionary potential for thermal adaptation [30,31], making our tropical *Drosophila*-parasitoid community a relevant system to study effects of future warming conditions on communities. *D. simulans* and parasitoid cultures were established between 2017 and 2018, identified using both morphology and DNA barcoding, and shipped to the Czech Republic under permit no. PWS2016-AU-002018 from Australian Government, Department of the Environment. All cultures were maintained at 23°C and 12:12 hour light and dark cycle at Biology Centre, Czech Academy of Sciences. The three larval parasitoid species *Asobara sp*. (Braconidae: Alysiinae; strain KHB, reference voucher no. USNMENT01557097, reference sequence BOLD process ID: DROP043-21), *Leptopilina sp*. (Figitidae: Eucolinae; strain 111F, reference voucher no. USNMENT01557117, reference sequence BOLD process ID: DROP053-21), and *Ganaspis sp*. (Figitidae: Eucolinae; strain 84BC, reference voucher no. USNMENT01557102 and USNMENT01557297, reference sequence BOLD process ID: DROP164-21) were used (for more details on the parasitoid strains see [32]). *Drosophila simulans* isofemale lines were kept on standard *Drosophila* medium (corn flour, yeast, sugar, agar, and methyl-4-hydroxybenzoate) for approximately 45 to 70 non-overlapping generations before the experiments. To revive genetic variation, five host lines were combined to establish two population cages of mass-bred lines before the start of the experiments. Single parasitoid isofemale lines were used and maintained for approximately 25 to 40 non-overlapping generations before the start of the experiment by providing them every week with two-day-old larvae of a different *Drosophila* species – *Drosophila melanogaster*.

### Experiments

We used a functional response approach following Mccoy’s framework to investigate the effects of warming on the strength of trophic and non-trophic interactions [26]. We first obtained each parasitoid functional response parameter at ambient and warmed temperatures with single-parasitoid treatments (Experiment 1). Then, we used these functional response parameter estimates to predict trophic interaction strength for each temperature and parasitoid combination with the null hypothesis that parasitoids were not interacting and thus had additive effects on host suppression. In Experiment 2 we empirically measured the effects of temperature and parasitoid combinations on trophic interaction strength and compared the predicted and observed values to identify emergent effects of multiple parasitoids on host suppression and their dependence on the temperature regime. We performed the two first blocks of Experiment 1 and entire Experiment 2 in parallel, and controls and single-parasitoid treatments were common to both experiments. In Experiment 3, we investigated the mechanisms of multiple parasitoid effects by dissecting hosts rather than rearing them. This allowed us to measure super- and multiparasitism rates and encapsulation depending on the temperature regime and parasitoid combinations.

A total of 22,920 *D. simulans* eggs were collected. 13,120 for experiment 1, 4,800 for experiment 2, and 5,000 for experiment 3 (from which 1,000 larvae were dissected). In experiments 1 and 2, 12,990 eggs (73%) successfully developed into adults (8,409 hosts and 4,581 parasitoids). The remaining 23% either died naturally or the host was suppressed without successful development of the parasitoid.

### Experiment 1: Single-parasitoid experiment

Eggs of *D. simulans* were placed in a single 90 mm high, and 28 mm diameter glass vial with 10mL of *Drosophila* media at six different densities replicated eight times each (5, 10, 15, 25, 50, or 100 eggs per 10mL of food media in a vial, total number of eggs = 13,120, N = 384 vials; Figure 1b). To collect *D. simulans* eggs, an egg-washing protocol was adapted from [33]. The day before the egg-washing protocol was conducted, two batches of egg-laying medium (Petri dishes with agar gel topped with yeast paste) were introduced in each population cage for flies to lay eggs overnight. Eggs were transferred to the experimental vials. We placed half of the vials at ambient temperature (22.7°C ± 0.4 s.d. - the current mean yearly temperature at the two study sites [28]), and the other half under elevated temperature (27.4°C ± 0.5 s.d. - projected change in global mean surface temperature for the late 21^st^ century is 3.7°C for the IPCC RCP8.5 baseline scenario [34]). Like in other *Drosophila* species, the thermal performance curve of *Drosophila simulans* demonstrates a decrease in performance from temperatures above 25°C [35].

**Figure 1.**
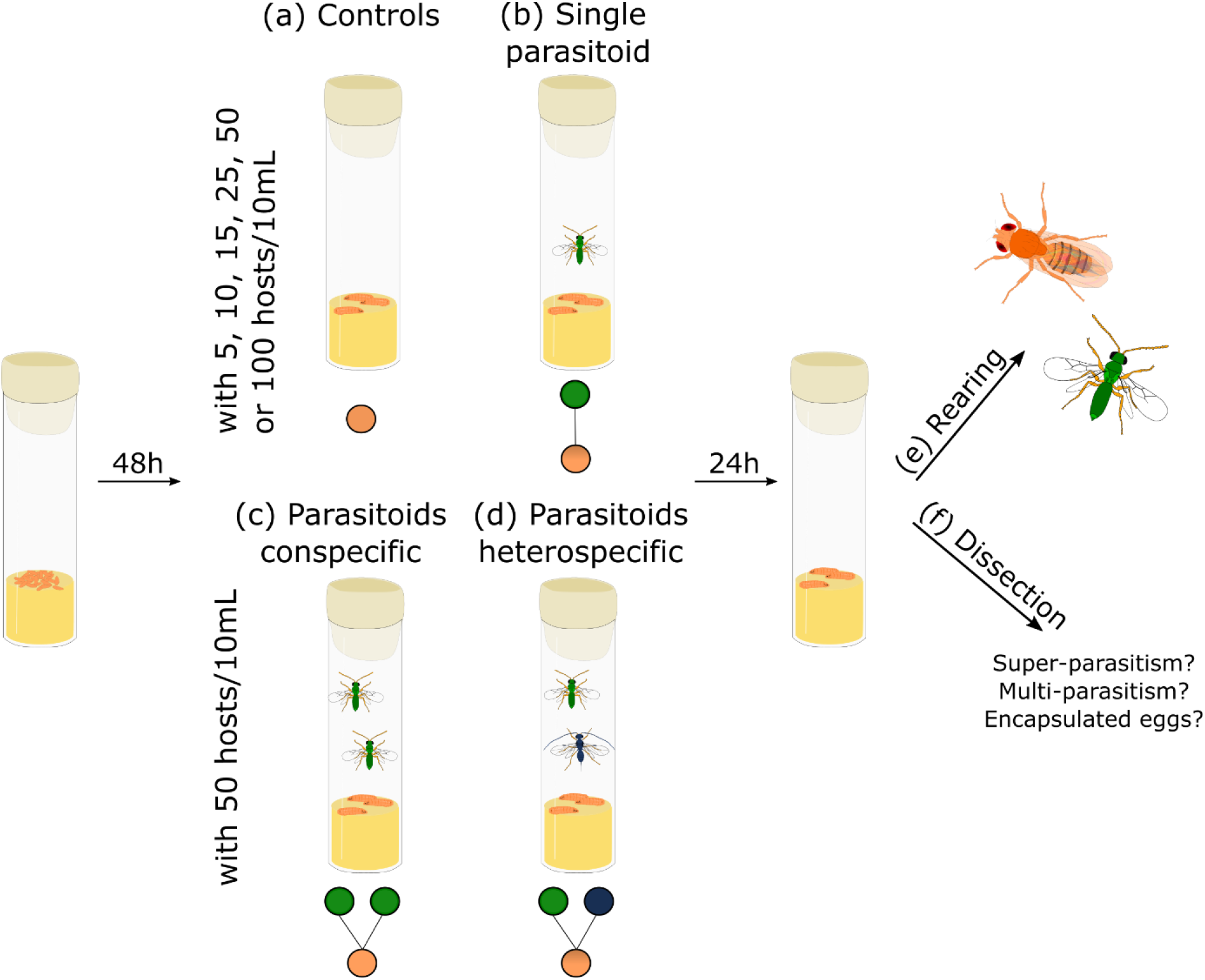
Schematic representation of the experimental design. (a) Controls or (b) one single parasitoid female with either 5, 10, 25, 50 or 100 *D. simulans* per 10 mL of media (N = 424 vials), (c) two parasitoids conspecific (N = 88 vials) or (d) two parasitoids heterospecific (N = 68 vials) with 50 *D. simulans* per 10 mL of media. (e) Rearing until adults emerge for Experiments 1 and 2 (up to 41 days), or (f) dissection of 10 3^rd^ instar larvae or pupae per vial two, three, or four days after infection for Experiment 3.

After 48 hours, we placed one single naïve mated three to five-day-old female parasitoid in each vial with *D. simulans* larvae. We removed the parasitoids twenty-four hours later. We repeated this for all three parasitoid species, temperatures, and host densities. We simultaneously performed controls without parasitoids to obtain the baseline for host survival without parasitism and potential variation during egg collection (Figure 1a). We checked vials were daily for adult emergences until the last emergence (up to 41 days for the species with the longest developmental time). We waited five consecutive days without any emergence to stop collecting and thus avoided confounding counts with a second generation. All emerged insects were collected, identified, sexed, and stored in 95% ethanol. Each treatment was replicated eight times across eight experimental blocks.

### Experiment 2: Multiple parasitoids experiment

To investigate the effect of warming on MPEs, we manipulated parasitoid assemblages and temperature in a fully factorial design (Figure 1c and d). We followed the same protocol described above for Experiment 1, using 50 *D. simulans* eggs per vial with two female parasitoids either from the same (Figure 1c) or different species (Figure 1d). Each treatment was replicated eight times across two blocks (N = 96). Controls and single-parasitoid treatments were standardized to experiment 1 with the 50 eggs per vial density. 50 eggs per standard *Drosophila* vial corresponds to low competition between hosts, a suitable host/parasitoid ratio when using one or two parasitoids, and enough host per vial for meaningful statistics.

### Experiment 3: Mechanisms of MPEs

In a follow-up experiment, we conducted a subset of the treatments described for Experiments 1 and 2 with *Asobara sp*. and *Ganaspis sp*. We put 50 *D. simulans* eggs per vial with 10 mL of food media under ambient and warming temperatures and introduced one parasitoid, two conspecific parasitoids, or the two heterospecific parasitoids, resulting in five different parasitoid assemblages. Each treatment was replicated ten times across two blocks (N = 100). Instead of rearing the insects to adults, we dissected ten 3^rd^ instar larvae or pupae per vial (Figure 1f). We individually transferred each host larva into a glass petri dish containing PBS and dissected it under a stereomicroscope. We recorded the number of parasitoid larvae and eggs of each species to assess superparasitism and multiparasitism events. When possible, we also identified the number and species of encapsulated parasitoids. Pictures of the eggs, larvae and encapsulated parasitoids for each species observed during the experiment are presented in Supplemental Material S1 Figure S1. At the elevated temperature, six replicates were dissected two days after infection (early dissection time) and four three days after infection (late dissection time). At the ambient temperature, four replicates were dissected three days after infection (early dissection time) and six four days after infection (late dissection time). We selected different times for dissection at each temperature to standardize parasitoid developmental stage while still being able to identify all the parasitoids that have parasitized the host. At the early dissection time, *Asobara sp*. were already at the larval stage, whereas *Ganaspis sp*. were still eggs. *Ganaspis* larvae were also observed at the late dissection time, sometimes simultaneously with a larva of *Asobara sp*. within the same host.

### Data analysis and modelling

#### Experiment 1: Single-parasitoid experiment

We combined numerical simulations of host density dynamics, accounting for host depletion [36]:

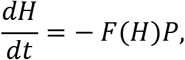

with Bayesian parameter estimation using the *rstan* package (e.g. [37]). *P* = 1 is the parasitoid density, and *F*(*H*) denotes the host density-dependent functional response. In the model fitting, we used Markov chain Monte Carlo to sample from the functional response’s model parameters’ posterior probability distribution *p*(*θ\H*_sup_) given the observations *H*_sup_, based on the likelihood function *p*(*H*_sup_|*θ*) and prior distributions *p*(*θ*), with *θ* the free parameters. *H*_sup_ is the number of *D. simulans* suppressed (the difference between adult hosts emerging from the controls without parasitoids and from the experiment). In each iteration, we computed numerical solutions of the equation with the built-in *Runge-Kutta* ODE solver to predict densities 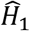 after one day for each given initial host density, *H*_0_. The initial host densities were taken from the average number of hosts that emerged from the controls for each density and temperature to account for potential deviations between the aimed and actual densities (Table S1). The likelihood was evaluated assuming a binomial distribution for observed numbers of suppressed hosts *H*_sup_ with *n* = *H*_0_ trials and 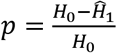 success probability. We used vague priors for all model parameters.

We fitted three different functional response models (Type II, Type III and generalized Type III) and retained the Type II functional response [38] after model comparison (see Supplement Material S2). The equation for the instantaneous attack rate of a parasitoid is as follows:

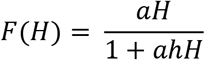

where *a* is the attack rate, and *h* is the handling time. Type II functional responses are thought to characterize the attack rate of many types of predators and parasitoids [39]. Parameter estimates and the functional responses for each species at each temperature are presented in Supplement Material S2 (Table S2 and Figure S2).

#### Experiment 2: Multiple parasitoids experiment

Host-parasitoid interaction strength was described with the combination of Degree of Infestation (DI; i.e., host suppression) and Successful Parasitism rate (SP; i.e., parasitoid performance). The observed degree of infestation (*DI_obs_*) and Successful parasitism rate (*SP*) were measured as:

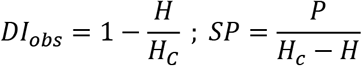

where *H* is the number of adult hosts emerging from the experiment vial, *H_C_* is the mean number of adult hosts emerging from the controls without parasitoids, and *P* is the number of parasitoid adults emerging from the experimental vial [40,41]. *DI_obs_* was set to zero if the number of hosts emerging from the treatment was greater than the controls. If no parasitoid emerged or the number of hosts suppressed was estimated to be zero, *SP* was set to zero. If the number of parasitoids that emerged was greater than the estimated number of hosts suppressed, *SP* was assigned to one. For treatments with parasitoid conspecifics, we assumed that each of the two parasitoid individuals was attacking the hosts equally; therefore, the number of parasitoid adults emerging was divided by two to calculate individual successful parasitism rate.

We analyzed these data with generalized linear models (GLMs) and verified model assumptions with the *DHARMa* package [42]. To correct for overdispersion of the residuals and zero inflation, data were modeled using zero-inflation models with a beta-binomial error distribution and a logit function using the *glmmTMB* function from the *TMB* package [43]. Two categories of predictor variables were used in separate models with temperature treatment (two levels: ambient and warming): (i) parasitoid treatment (three levels; single parasitoid, two parasitoids conspecific, and two parasitoids heterospecific), and (ii) parasitoid species assemblage (nine levels). The two-way interactions between temperature and either parasitoid treatment or parasitoid assemblage were tested and kept in our models if judged to be significant based on backward model selection using Likelihood-ratio tests. The significance of the effects was tested using Wald type III analysis of deviance with Likelihood-ratio tests. Factor levels were compared using Tukey’s HSD *post hoc* comparisons of all means and the *emmeans* package [44]. Results for developmental rate are presented in Supplement Material S3 Figure S3.

#### Estimation of multiple parasitoid effects

To predict the degree of infestation if parasitoids have independent effects on host suppression, we used the method developed by Mccoy *et al*. [26], which considers host depletion. This method uses the functional responses obtained from Experiment 1 in a population-dynamic model to predict how host density changes in time as a function of initial density and parasitoid combination for each temperature. We thus calculated the estimated Degree of Infestation (*DI_0_*) by integrating the aggregate attack rates over the duration of the experiment as host density declines. We first solved the equation

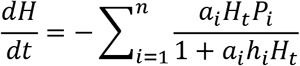

similar to the equation described for Experiment 1 but adapted to *n* parasitoids. Then we calculated the estimated Degree of Infestation as

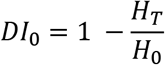

where *H_0_* is the initial host density, and *H_T_* is the estimated host population at the end of the experiment (time *T* = 1 day). This method allows a reasonable estimate of *DI_0_* for the null hypothesis that predators do not interact [27]. The lower and upper confidence intervals (CI) around the predicted values were estimated with a global sensitivity analysis based on the functional response parameters estimates to generate 100 random parameter sets using a Latin hypercube sampling algorithm [45]. The expected degree of infestation was calculated for each parameter set using the *sensRange* function in the R package *FME*. The 2.5% and the 97.5% quantiles of the values obtained from these simulations were used as 95% CIs around the predictions.

Predictions from the population dynamic model were then compared with the observed values (*DI_obs_*). Estimated DI values greater than observed DI translate to risk reduction, while lower estimates reflect risk enhancement for the host with multiple parasitoids. We calculated the difference between *DI_obs_* and mean *DI_0_* for each treatment and investigated the effects of temperature (ambient versus warmed), parasitoid diversity (one or two species), and their interaction if significant, using an analysis of variance (ANOVA) with the *aov* function. We statistically compared the observed and estimated DI for each temperature regime using GLMs with a beta-binomial error distribution and a logit function with *DI_0_* as an offset (i.e., predictor variable) following Sentis et al. (2017). A positive or negative significant intercept indicates that *DI_0_* values underestimate or overestimate *DI_obs_*, respectively.

#### Experiment 3: MPEs mechanisms

The frequency of super- and multiparasitism events was calculated out of the larvae parasitized per vial (total of 1,000 larvae dissected across 100 vials, out of which 868 were parasitized: presence of either one or both parasitoid species and/or trace of melanization). The frequency of encapsulated parasitoids was calculated out of the total number of parasitoids per larva. Effects of temperature and parasitoid assemblages on these frequencies were analyzed with generalized linear mixed models (GLMMs) with the method described for Experiment 2 and the time of dissection (early or late) as a random effect. All analyses were performed using R 4.0.2 [46].

## Results

### Effects of multiple parasitoids on host suppression under warming

The degree of infestation observed in the experiment varied from the model estimations (Figure 2). Temperature significantly affected these differences (F_1,93_ = 13.9, P < 0.0001), but parasitoid diversity did not (F_1,93_ = 0.09, P = 0.766) (Table S3, Table S4), implying that parasitoid density rather than their diversity is important for host suppression. The comparison of the estimated and observed DI revealed that, in most cases, there was no significant difference between predicted and observed DI at ambient temperature, implying neutral effects with multiple parasitoids (when looking at the intercept of the GLM with *DI*_*0* as_ an offset; value ± SE: 0.12 ± 0.32, z value = 0.381, df = 40, P = 0.703), whereas under warming the predicted *DI_0_* significantly underestimated the observed *DI_ob_*, implying risk enhancement for the host (value ± SE: 0.44 ± 0.18, z value = 2.431, df = 40, P = 0.015; Figure 2).

**Figure 2.**
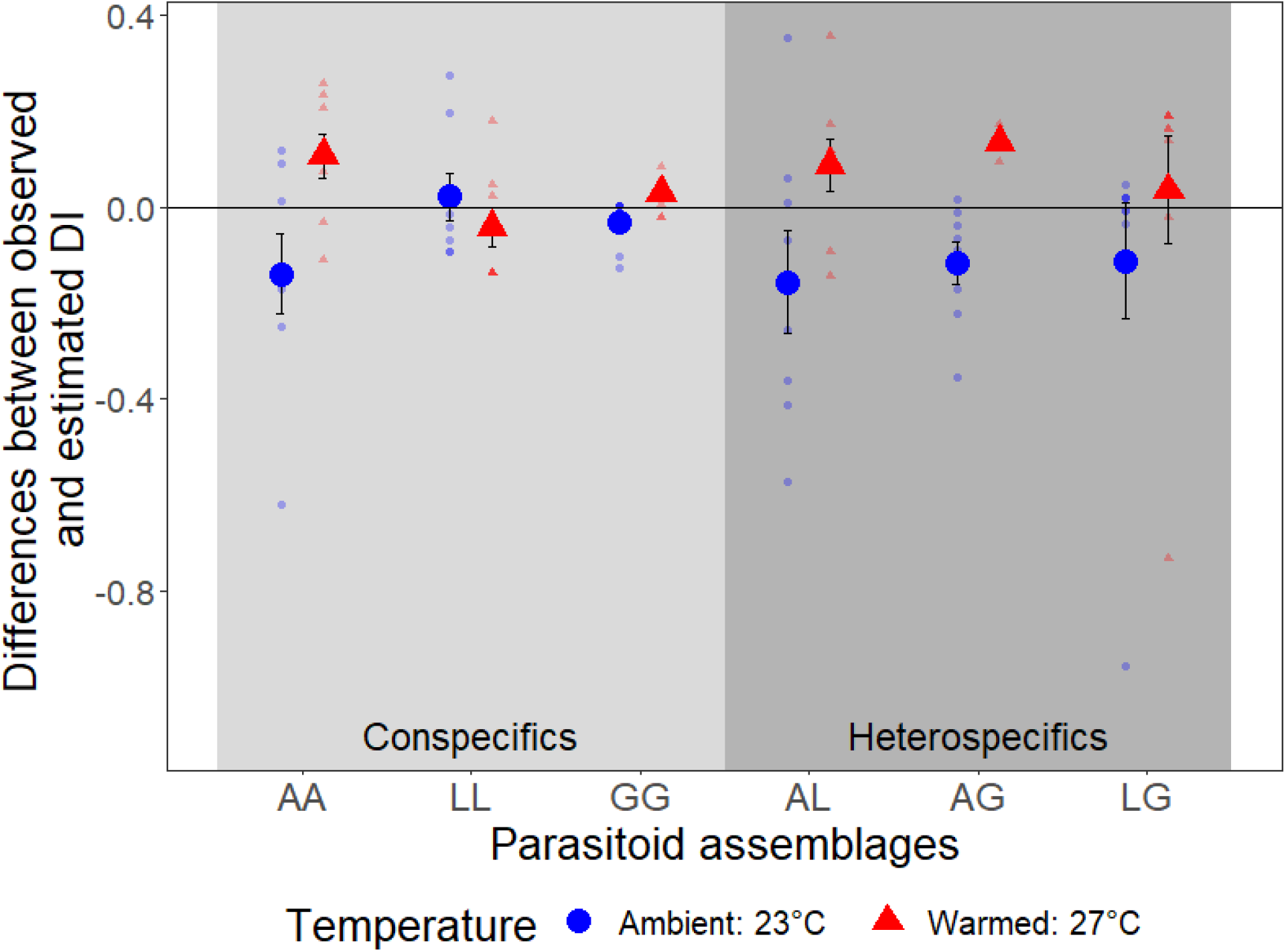
Differences between observed and estimated degree of infestation (DI) for each parasitoid assemblage and temperature (i.e., *DI_obs_* - *DI_0_*). Negative values translate to risk reduction, while positive values reflect risk enhancement for the host with multiple parasitoids. Light grey panel: two parasitoids conspecific, darker grey panel; two parasitoids heterospecific. Parasitoid abbreviations: A: *Asobara sp*., L: *Leptopilina sp*., and G: *Ganaspis sp*. Big dots represent the means (±SE), and small dots represent raw data.

### Effects of warming and parasitoid assemblages on the observed degree of infestation

Contrary to the effects of multiple parasitoids on host suppression, the observed degree of infestation *DI_obs_* was not significantly affected by temperature (χ^2^_(1)_ = 1.05, P = 0.306), or parasitoid treatment (single, two conspecific or two heterospecific parasitoid assemblages: χ^2^_(2)_ = 4.26, P = 0.119; Table S5) due to species-specific effects. DI only varied with parasitoid species assemblages (χ^2^_(8)_ = 251.92, P < 0.0001, Table S6). DI was the highest in assemblages with *Ganaspis sp*., either alone, with a conspecific, or another parasitoid species (Figure S4).

### Effect of warming and parasitoid assemblages on parasitoid performance

Despite having no effect on DI, parasitoid treatment (single, two conspecific, or two heterospecific parasitoid assemblages) significantly affected successful parasitism rate, and the effect varied among parasitoid species (two-way interaction: χ^2^_(4)_ = 16.88, P = 0.002; Table 1; Table S7). SP of *Ganaspis sp*. decreased by 95.7% (95% CI: 93.6 - 97.8%) with the presence of a parasitoid conspecific [*Post hoc* Odds Ratio (OR) = 0.043, P < 0.0001 for contrast 2Pc/1P; Table S8], and by 83.4% (CI: 75.4 - 91.3%) with the presence of a parasitoid heterospecific compared to when alone (OR = 0.166, P < 0.001 for contrast 2Ph/1P; Table S8). However, it increased by 287.6% (CI: 178.8 - 396.4%) when the parasitoid competitor was from another species compared to a conspecific (OR = 3.876, P < 0.0001, for contrast 2Pc/2Ph; Table S8). SP of *Asobara sp*. decreased by 55.2% (CI: 41.5 - 69.7%) when a parasitoid conspecific was present compared to when alone (OR = 0.448, P = 0.036), but was not significantly affected by the presence of a parasitoid heterospecific (OR = 0.712, P = 0.484). There were no significant effects of parasitoid treatments for SP of *Leptopilina sp*. Effects of parasitoid assemblages on SP also varied between parasitoid species and are presented in Supplementary Material S6 (Table S9, Table S10, and Figure S5).

**Table 1.** Odds ratios of a successful parasitism event between parasitoid treatments (single parasitoid, two parasitoids conspecific, and two parasitoids heterospecific) for each parasitoid species. Results are averaged over both temperatures because there was no significant interaction between temperature and parasitoid treatments. Values less than or greater than one denote a decrease or an increase in the odds of successful parasitism, respectively. Significant differences are highlighted in bold.

| Parasitoid species | Contrast | Odds Ratio | P-value |
| --- | --- | --- | --- |
| *Ganaspis sp.* | 2 conspecifics/single | **0.043** | **<0.0001** |
|  | 2 heterospecifics/single | **0.166** | **0.0007** |
|  | heterospecifics/conspecifics | **3.876** | **<0.0001** |
| *Asobara sp.* | 2 conspecifics/single | **0.448** | **0.036** |
|  | 2 heterospecifics/single | 0.711 | 0.484 |
|  | heterospecifics/conspecifics | 1.589 | 0.251 |
| *Leptopilina sp.* | 2 conspecifics/single | 0.182 | 0.494 |
|  | 2 heterospecifics/single | 0.871 | 0.994 |
|  | heterospecifics/conspecifics | 4.764 | 0.295 |

Effects of temperature on SP also depended on the species (two-way interaction: χ^2^_(2)_ = 7.31, P = 0.026; Table S7). Only *Ganaspis sp*. was significantly affected by temperature, and its SP decreased by 58.8% (CI: 69.8 - 47.8%) with warming (OR = 0.412, χ^2^_(1)_ = 10.17, P = 0.001). However, all species developed faster under warming (Figure S3).

### Mechanisms of MPEs

The frequency of either super- or multiparasitism events, reflecting strength of intrinsic competition among parasitoids, was significantly affected by parasitoid assemblages (χ^2^_(4)_ = 572.40, P < 0.0001), temperature (χ^2^_(1)_ = 4.49, P = 0.034), and the interaction between parasitoid assemblages and temperature χ^2^_(4)_ = 36.04, P < 0.0001; Figure 3, Table S11). Superparasitism rate increased by 239% (CI: 230-308%) when *Ganaspis sp*. was with a conspecific (OR = 3.69, P < 0.0001), and by 581% (CI: 411-751%) when *Asobara sp*. was with a conspecific (OR = 6.81, P < 0.0001) compared to when they were alone, but without significant differences between temperature treatments. In the parasitoids heterospecific treatments, warming significantly increased the frequency of super- and multiparasitism events by 173% (CI: 130-216%; OR = 2.73, P < 0.0001), indicating an increase in intrinsic competition among parasitoids with warming.

**Figure 3.**
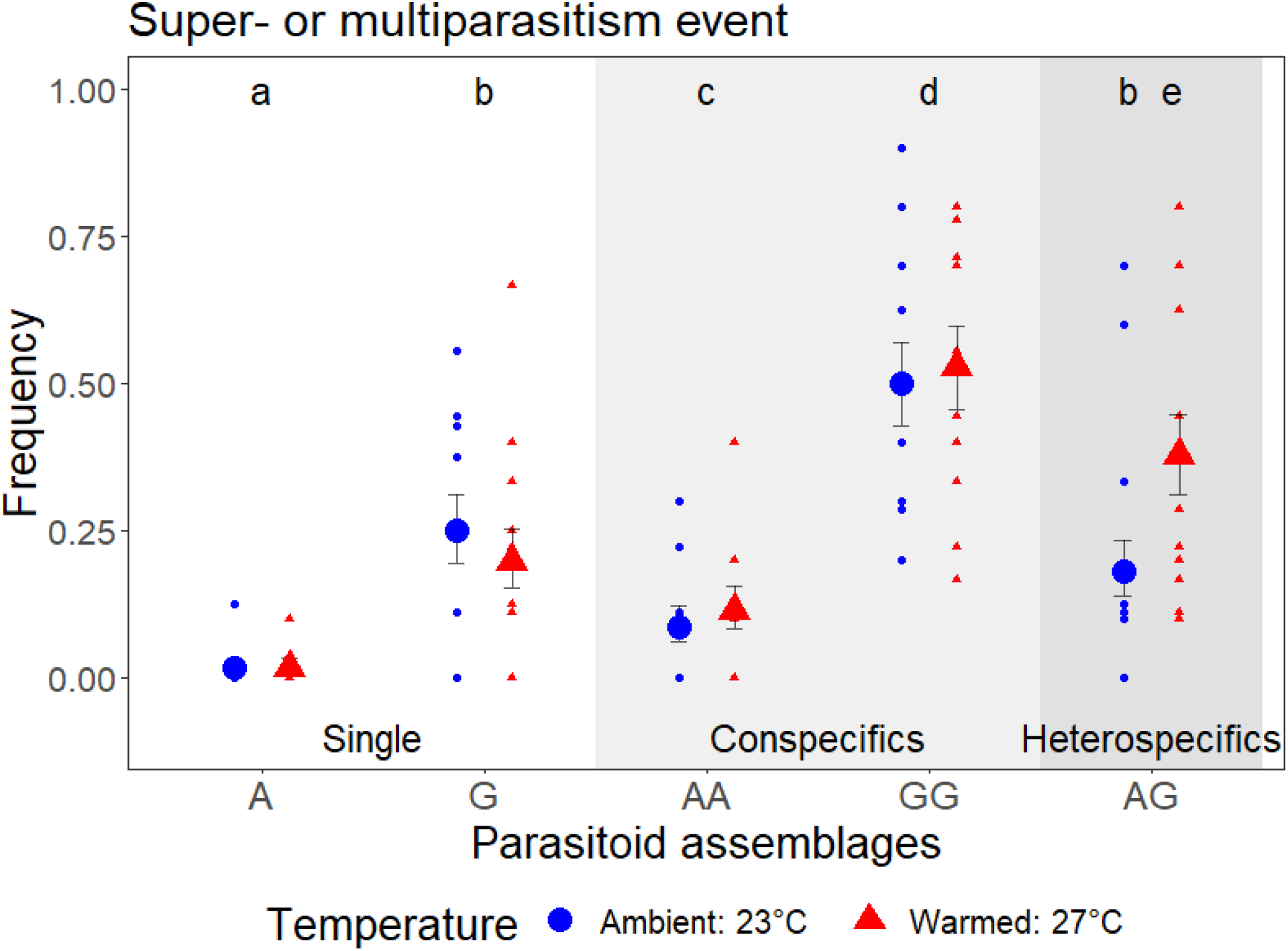
Frequency of super- or multiparasitism events out of the total of parasitized hosts per vial (N = 100 vials) significantly changed depending on parasitoid assemblage and temperature regime, indicating changes in intrinsic competitive interaction strength among parasitoids. Within each plot, different small letters denote significant differences between parasitoid assemblages (and temperature regime if significant). White panel: single parasitoid, light grey panel: two parasitoids conspecific, darker grey panel; two parasitoids heterospecific. Parasitoid abbreviations: A: *Asobara sp*., and G: *Ganaspis sp*. Big dots represent the estimated means (±95% CIs), and small dots represent raw data.

The frequency of encapsulated parasitoids differed between parasitoid species, but not between treatments (results presented in Supplement Material S8: Table S12 and Figure S6), indicating that host immune response did not change depending on the treatments.

## Discussion

The key result from our study is the synergistic effects of multiple predators for top-down control at elevated temperature across predator assemblages. However, parasitoid performance often decreased when multiple parasitoids were present due to intrinsic competition among parasitoids, potentially limiting the long-term benefits for ecosystem functioning.

### Warming increases the effects of multiple predators on the risk of predation

Our results showed that warming led to a higher top-down control than expected with multiple predators. Indeed, our mathematical model underestimated trophic interaction strength measured in multiple predators treatments at elevated temperatures. Our results are in concordance with previous studies on diverse systems on the importance of considering non-trophic interactions to predict the effect of multiple predators on top-down control under global changes. Drieu et al. [47] found that predator diversity enhanced the biological control of insect pests in vineyards under warming due to functional complementarity among predator species, while effects were substitutive at ambient temperature. Cuthbert et al. [11] also found an impact of temperature on intraspecific multiple predator effects on an invasive Gammaridae species (Amphipoda). Yet, the direction of the effects contrasted ours as they observed risk enhancement at low temperature and risk reduction with warming. Sentis et al. [10] found a general trend of predation risk reduction for the prey with multiple predators in an aquatic food web but without any effect of temperature on those emergent MPEs. Our study goes further by showing the important impact of warming on the impacts of multiple predators on prey suppression across multiple assemblages of conspecifics and heterospecifics. In addition to increasing prey suppression with multiple predators under warming in terrestrial ecosystems, a diverse predator community also increases the chances of complementarity in the face of environmental variation and disturbance [48]. Indeed, the presence of multiple predator species could mitigate the adverse effects of warming on top-down control due to resource partitioning and/or functional redundancy [47,49,50]. Therefore, preserving predator biodiversity should be generally beneficial for top-down control under climate change.

### Mechanisms behind emergent multiple predator effects on the prey

Because of the synergistic effects of multiple parasitoids on host suppression under warming found in our study, we could have hypothesized that warming weakens interference between parasitoids, similarly to predator-prey systems [51]. However, our host-parasitoid system allowed us to investigate further the potential mechanisms behind our results, especially the strength of intrinsic competitive interactions between parasitoids (i.e., frequency of superparasitism and multiparasitism events). We found generally higher intrinsic competition in multiple parasitoid treatments than single parasitoid treatments and higher intrinsic competition under warming when the two species were present compared to ambient temperature. When superparasitized or multiparasitized, the host was less likely to survive, possibly because its immune response was less likely to overcome multiple parasitoids. Therefore, the higher top-down control observed under warming with multiple parasitoids was due to a higher parasitism pressure and not because of weaker interactions between parasitoids.

We conducted the experiments in simplified laboratory conditions and forced parasitoids to share the same habitat (a vial) and overlapped in time (24 hours), which does not allow for resource partitioning [52]. This might have enhanced the rate of super- and multi-parasitism events and thus top-down control. In nature, warming could also change predator habitat use [8,9], and phenology [53,54], leading to changes in MPEs. However, the impact of temperature on MPEs was consistent across parasitoid assemblages, suggesting a general pattern for synergistic effects with multiple natural enemies under warming in our system.

### Parasitoid performance was mostly affected by parasitoid assemblage

Despite multiple parasitoids enhancing host suppression under warming, the successful parasitism rate was often lower at both temperatures when another parasitoid individual was present, probably due to the strong intrinsic competitive interactions observed through dissections. A decrease in parasitoid performance would potentially limit the synergistic effects of multiple parasitoids for host suppression in the long term. Similarly, another study on *Drosophila-parasitoid* interactions observed a significant impact of thermal regime on parasitoid success, but still without changes in the observed degree of infestation [55]. The long-term effects of warming on parasitoid populations are thus uncertain, and hosts from the next generation might benefit from lower parasitoid abundances due to a lower rate of successful parasitism.

### Similar effects of intra-versus interspecific multiple parasitoids on top-down control

Similar to other studies, we did not find significant differences between treatments with multiple conspecifics or heterospecific predators for prey suppression [56–58]. Therefore, it is essential to look at the effects of both predator diversity and density on prey suppression, rather than only using a substitutive approach (i.e., keeping predator density constant [52]), which might confound the results. When niche differentiation is allowed, for example, with habitat heterogeneity or a more prolonged timeframe that includes potential differences in phenology, an increase in predator diversity should intensify prey suppression because of functional diversity rather than because of diversity *per se* [58–60]. Here, two predators of the same species rather than a single predator intensified prey suppression at warmer temperatures despite the small scale of the experiment. Allowing for differentiation in habitat domain between predator species might have yielded higher prey suppression in treatments with heterospecifics and a lower rate of multiparasitism. Given the likely ubiquity of resource partitioning in nature [61], preserving predator biodiversity would be the best strategy to maintain top-down control.

### No effects of treatments on the observed degree of infestation

Warming had a significant effect on the differences between observed and estimated degree of infestation. However, temperature treatment had no significant effect on the observed degree of infestation. Moreover, prey suppression was generally higher when predator assemblages included the best-performing species, *Ganaspis sp*., no matter the predator treatment or temperature. A meta-analysis on the effects of predator diversity on prey suppression found a similar trend across the 46 studies taken into account [62], but also found a general positive effect of multiple predators on top-down control. Contrastingly, a meta-analysis of 108 biological control projects found no relationship between the number of agents released and biological control success for insect pests [63]. However, increasing predator diversity should be generally beneficial for top-down control by increasing the chances to have a more effective natural enemy species in the community, as was the case in our study (i.e., sampling effect model [64]). Moreover, the presence of multiple species in the community could buffer any mismatch between predator and prey species induced by warming [65]. *Ganaspis sp*. was the best performing species in suppressing *D. simulans* across treatments. Still, its performance decreased with warming, suggesting that parasitism rate, and therefore host suppression, could also be reduced in the longer term due to a decrease in parasitoid population.

### Conclusion

Overall, pairwise interaction strength generally failed to accurately estimate the trophic interaction strength observed, indicating that non-trophic interactions must be considered to predict the effects of multiple predators on prey suppression and in food web studies in general [66]. While previous studies show altered MPEs with warming due to changes in resource partitioning [8,11], our study is the first, to our knowledge, to show signs of direct effects of warming on predator interactions across predator assemblages, resulting in a higher top-down control with multiple predators at elevated temperature.

## Acknowledgements

We thank Anna Mácová, Andrea Weberova, Grégoire Proudhom, and Joel Brown for their help during the setup of the experiments, and Tereza Holicová and Vincent Montbel for their help dissecting the larvae. Tereza Holicová made the drawings used for Figure 1. We acknowledge funding support from the Czech Science Foundation grant no. 20-30690S. BR acknowledges the support of iDiv funded by the German Research Foundation (DFG–FZT 118, 202548816).

