## Supplement Materials for "Presence of multiple parasitoids decreases host survival under warming, but parasitoid performance also decreases"

*Supplement Material S1: Identification of parasitoid eggs and larvae inside D. simulans 2, 3, or 4 days after infection.*

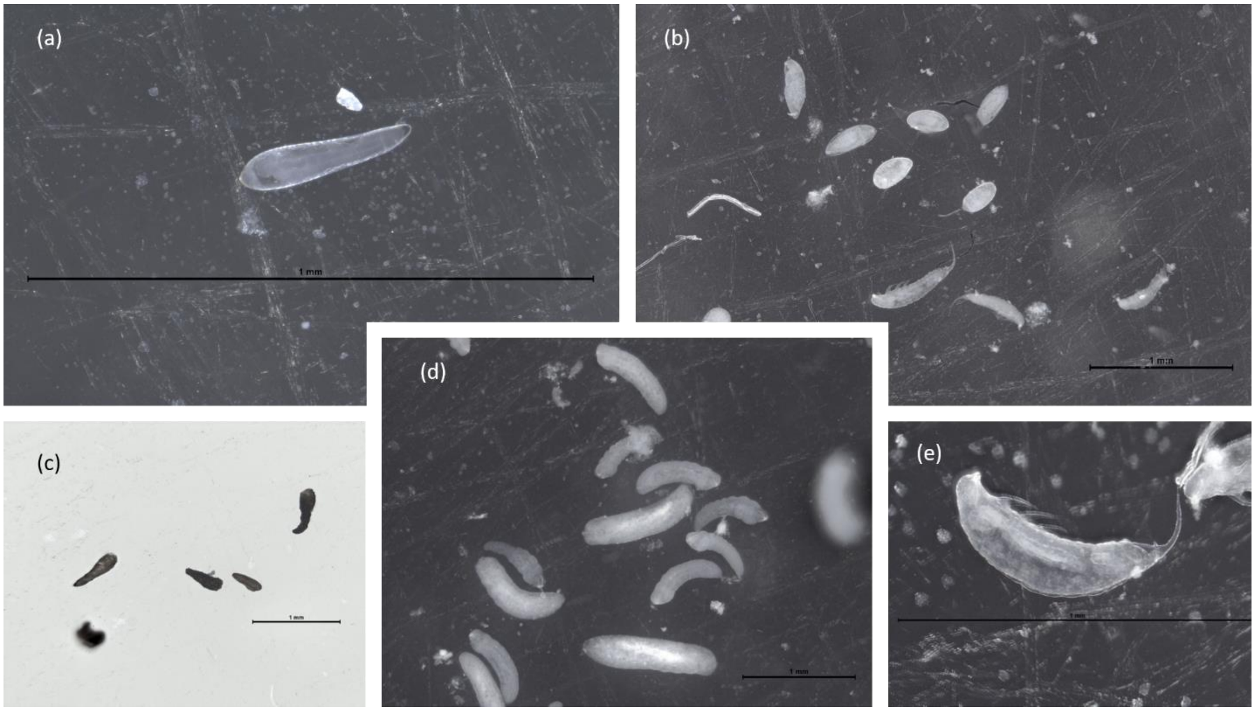

**Figure S1.** (a) *Asobara sp*. egg, (b) *Ganaspis sp*. eggs, and larvae, (c) encapsulated parasitoid eggs, (d) *Asobara sp*. larvae, (e) *Ganaspis sp*. larva. For all pictures, the bar scale represents 1 mm

*Supplement Material S2: Parasitoid functional responses*

We fit three functional response models to the single-parasitoid experiments (Experiment 1) at all temperatures and parasitoid species. All three functional response models can be expressed by

$$F(H) = \frac{aH^{1+q}}{1 + ahH^{1+q}}$$

where (1) $q$ = 0 defines a type II response, (2) $q$ = 1 defines a type III response, and (3) a free parameter $q$ defines a generalized type III response that allows a continuous shift between type II and type III (Rosenbaum & Rall, 2018). We used the leave-one-out information criterion (LOOIC) for model comparison, which was computed from the log-likelihood values of posterior samples (*loo* package). Although type III and generalized type III responses had lower LOOIC scores than the type II response (differences ΔLOOIC = 0.7, SE = 30.6, and ΔLOOIC = 19.2, SE = 26.2, respectively), the differences were in the range of estimated uncertainty. Therefore, we chose the type II response as the most parsimonious model.

**Table S1.** Actual initial host density based on mean host number emerging from the controls ± standard error.

| **Temperature** | **Aimed host density** | **Actual host density ± s.e.** |
|---|---|---|
| 23°C | 5 | 5 ± 0.6 |
| 23°C | 10 | 9 ± 0.5 |
| 23°C | 15 | 13 ± 0.5 |
| 23°C | 25 | 21 ± 0.7 |
| 23°C | 50 | 38 ± 4.2 |
| 23°C | 100 | 83 ± 1.7 |
| 27°C | 5 | 4 ± 0.3 |
| 27°C | 10 | 6 ± 0.5 |
| 27°C | 15 | 12 ± 1.5 |
| 27°C | 25 | 20 ± 0.9 |
| 27°C | 50 | 38 ± 5.3 |
| 27°C | 100 | 79 ± 5.2 |

**Table S2.** Estimated parameters *a* search rate (day host$^{-1}$) and *h* handling time (day host$^{-1}$) of the type II functional response for each parasitoid species at each temperature ± standard error.

| Species | Temperature | a ± s.e. | h ± s.e. |
|---|---|---|---|
| *Asobara sp.* | 23°C | 1.85 ± 0.16 | 0.029 ± 0.002 |
| *Asobara sp.* | 27°C | 0.56 ± 0.05 | 0.008 ± 0.003 |
| *Ganaspis sp.* | 23°C | 3.13 ± 0.21 | 0.002 ± 0.001 |
| *Ganaspis sp.* | 27°C | 1.26 ± 0.05 | 0.001 ± 0.0004 |
| *Leptopilina sp.* | 23°C | 1.67 ± 0.58 | 0.541 ± 0.064 |
| *Leptopilina sp.* | 27°C | 0.08 ± 0.01 | 0.042 ± 0.026 |

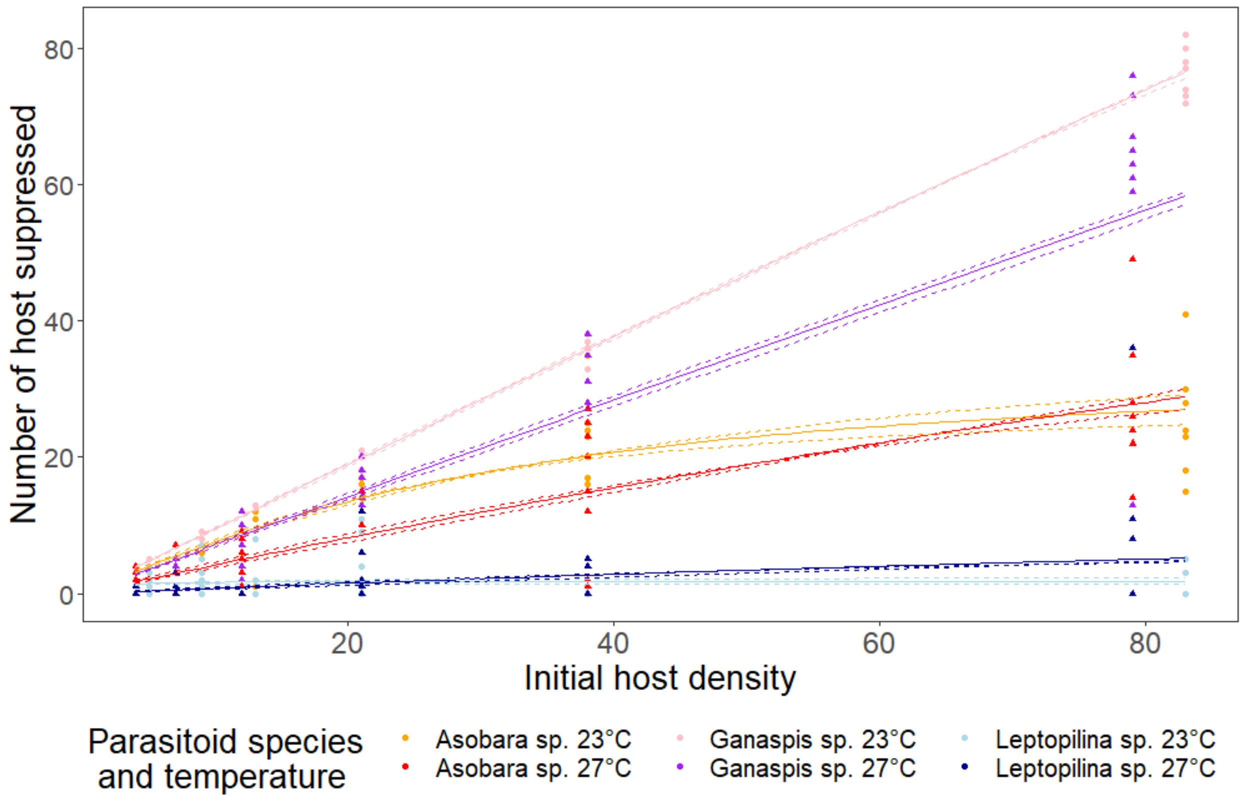


**Figure S2.** Type II functional responses of the three parasitoids at ambient (23°C) and warmed (27°C) temperature estimated from Experiment 1 (N = 288 vials). Points represent observed values, solid lines correspond to the fitted functional responses and dashed lines the 95% confidence intervals

*Supplement Material S3: Effects of warming and parasitoid assemblage on parasitoid developmental rate*

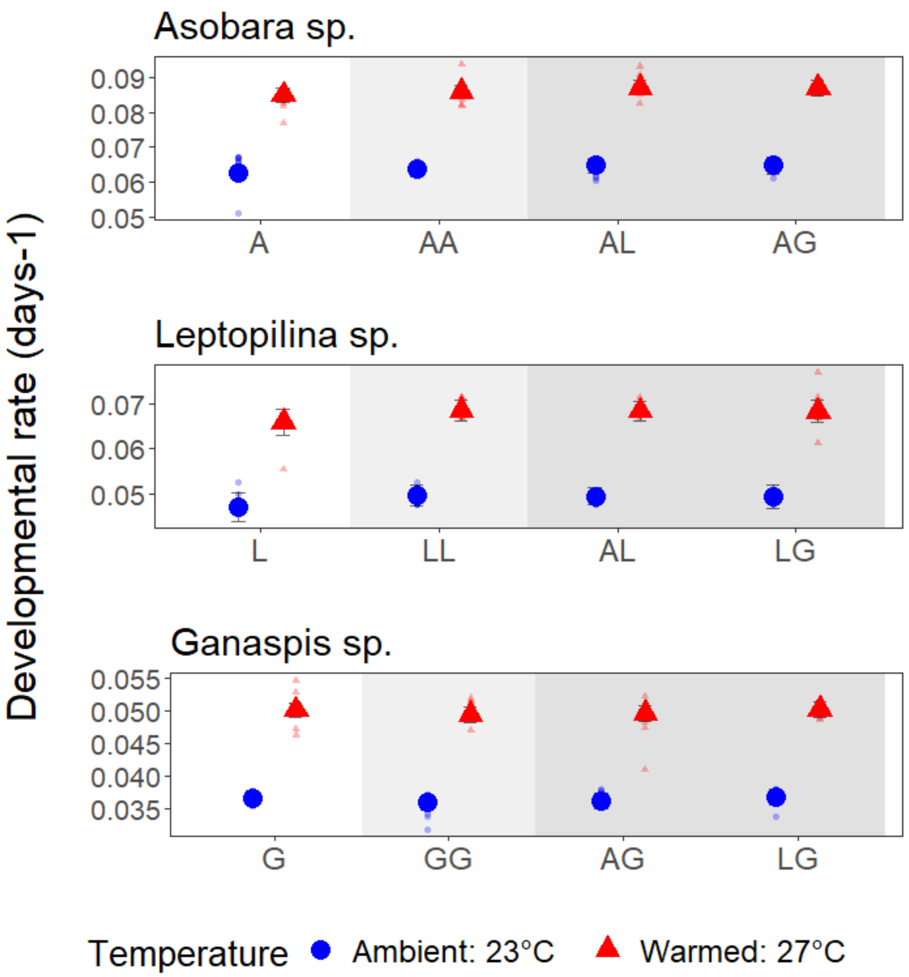


**Figure S3.** Development rate per day of each parasitoid species significantly increased with warming but was not affected by parasitoid assemblage. White panel: single parasitoid, light grey panel: two parasitoids conspecific, darker grey panel; two parasitoids heterospecific. Parasitoid abbreviations: A: *Asobara sp*., L: *Leptopilina sp*., and G: *Ganaspis sp*. Big dots represent the estimated means (±95% CIs), and small dots represent raw data. Note that the y-axis scale varies between parasitoid species.

*Supplement Material S4: Effect of warming and parasitoid assemblages on the differences between observed and estimated degree of infestation*

**Table S3.** Effects of temperature and parasitoid treatment on the differences between observed and estimated degree of infestation. Effects are shown by the Anova table (Type III test) with the effects of temperature (2 levels) and parasitoid treatment (3 levels: single parasitoid, two conspecifics, two heterospecifics). Degrees of freedom (Df) are given for each factor and the residuals.

| **Effects** | **F-value** | **Df** | **p-value** |
|---|---|---|---|
| Temperature | 13.9 | 1 | < 0.0001 |
| Parasitoid treatment | 0.09 | 1 | 0.438 |
| | | 93 | |

**Table S4.** Effects of temperature and parasitoid assemblage on the differences between observed and estimated degree of infestation. Effects are shown by the Anova table (Type III test) with the effects of temperature (2 levels) and parasitoid assemblage (9 levels). Degrees of freedom (Df) are given for each factor and the residuals.

| **Effects** | **F-value** | **Df** | **p-value** |
|---|---|---|---|
| Temperature | 13.4 | 1 | < 0.0001 |
| Parasitoid assemblage | 0.14 | 5 | 0.982 |
| | | 89 | |

*Supplement Material S5: Effect of warming and parasitoid assemblages on the degree of infestation*

**Table S5.** Effects of temperature and parasitoid treatment on degree of infestation. Effects are shown by the summary of Likelihood-ratio chi-square tests with the effects of temperature (2 levels) and parasitoid treatment (3 levels: single parasitoid, two conspecifics, two heterospecifics). Degrees of freedom (Df) are given for each factor and the residuals.

| **Effects** | **$\chi$2** | **Df** | **p-value** |
|---|---|---|---|
| Temperature | 1.05 | 1 | 0.306 |
| Parasitoid treatment | 4.26 | 2 | 0.119 |
| | | 138 | |

**Table S6.** Effects of temperature and parasitoid assemblage on the degree of infestation. Effects are shown by the summary of Likelihood-ratio chi-square tests with the effects of temperature (2 levels) and parasitoid assemblage (9 levels). Degrees of freedom (Df) are given for each factor and the residuals.

| **Effects** | **$\chi$2** | **Df** | **p-value** |
|---|---|---|---|
| Temperature | 3.42 | 1 | 0.064 |
| Parasitoid assemblage | 251.92 | 8 | < 0.0001 |
| | | 132 | |

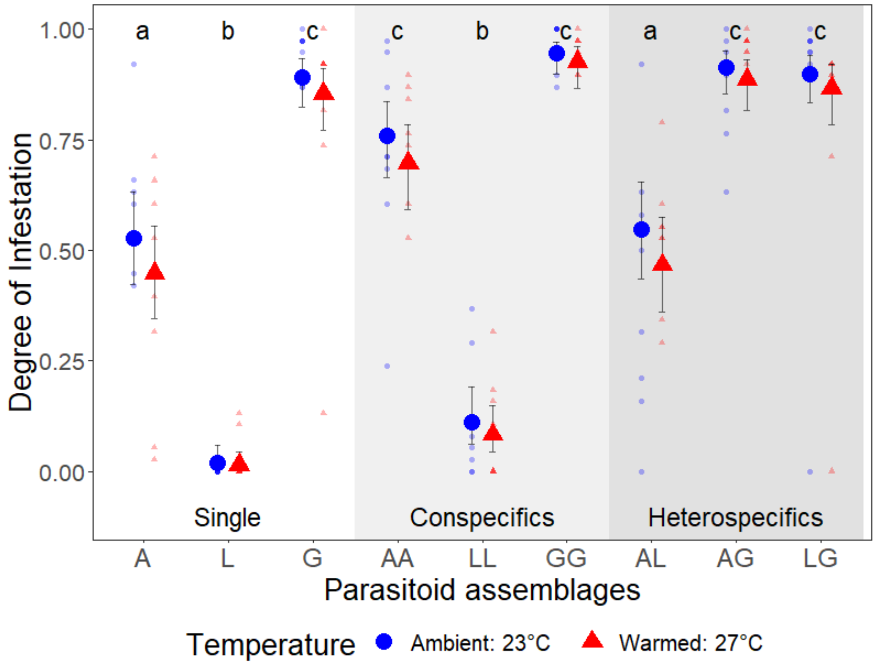


**Figure S4.** Degree of infestation for each parasitoid assemblage and temperature (N = 144 vials). Different small letters denote significant differences between parasitoid assemblages. White panel: single parasitoid, light grey panel: two parasitoids conspecific, darker grey panel; two parasitoids heterospecific. Parasitoid abbreviations: A: *Asobara sp*., L: *Leptopilina sp*., and G: *Ganaspis sp*. Big dots represent the estimated means (±95% CIs), and small dots represent raw data.

*Supplement Material S6: Effects of warming and parasitoid assemblage on successful parasitism rate*

**Table S7.** Effects of temperature, parasitoid treatment, parasitoid species, and their interactions on successful parasitoid rate. Effects are shown by the summary of Likelihood-ratio chi-square tests with the effects of temperature (2 levels), parasitoid treatment (3 levels: single parasitoid, two conspecifics, two heterospecifics), and parasitoid species (3 levels: *Asobara sp*., *Leptopilina sp*., and *Ganaspis sp*.). Degrees of freedom (Df) are given for each factor and the residuals.

| Effects | $\chi^2$ | Df | p-value |
|---|---|---|---|
| Temperature | 2.17 | 1 | 0.140 |
| Parasitoid treatment | 13.38 | 2 | 0.001 |
| Parasitoid sp. (PS) | 96.87 | 2 | < 0.0001 |
| Temperature x PS | 7.31 | 2 | 0.026 |
| Treatment x PS | 16.88 | 4 | 0.002 |
| | | 178 | |

**Table S8.** Effects of parasitoid treatment on successful parasitism rate for each parasitoid species. Abbreviations: 1P: single parasitoid, 2Pc: two parasitoids conspecific, and 2Ph: two parasitoids heterospecific. Results are averaged over both temperatures because there was no significant interaction between temperature treatments and parasitoid treatment.

| Parasitoid species | Contrast | Odds Ratio | p-value |
|---|---|---|---|
| *Asobara sp.* | 2Pc/1P | 0.45 | 0.0363 |
| | 2Ph/1P | 0.71 | 0.484 |
| | 2Pc/2Ph | 1.59 | 0.251 |
| *Ganaspis sp.* | 2Pc/1P | 0.04 | < 0.0001 |
| | 2Ph/1P | 0.17 | 0.0007 |
| | 2Pc/2Ph | 3.88 | < 0.0001 |
| *Leptopilina sp.* | 2Pc/1P | 0.18 | 0.494 |
| | 2Ph/1P | 0.87 | 0.99 |
| | 2Pc/2Ph | 4.76 | 0.30 |

**Table S9.** Effects of temperature and parasitoid assemblage on successful parasitoid rate for each parasitoid species. Effects are shown by the summary of Likelihood-ratio chi-square tests with the effects of temperature (2 levels) and parasitoid assemblage (4 levels). Degrees of freedom (Df) are given for each factor and the residuals.

| Parasitoid species | Effects | $\chi 2$ | Df | p-value |
|---|---|---|---|---|
| *Asobara sp.* | Temperature | 0.013 | 1 | 0.909 |
| | Para. assemblage | 16.48 | 3 | < 0.0001 |
| | | | 57 | |
| *Leptopilina sp.* | Temperature | 0.080 | 1 | 0.777 |
| | Para. assemblage | 7.55 | 3 | 0.056 |
| | | | 57 | |
| *Ganaspis sp.* | Temperature | 10.17 | 1 | 0.001 |
| | Para. assemblage | 46.66 | 3 | < 0.0001 |
| | | | 57 | |

**Table S10.** Effects of parasitoid assemblages on successful parasitism rate for each parasitoid species. Abbreviations: A: *Asobara sp.*, L: *Leptopilina sp.*, and G: *Ganaspis sp.* Results are averaged over both temperatures because there was no significant interaction between temperature treatments and parasitoid assemblages.

| Parasitoid species | Contrast | Odds Ratio | p-value |
|---|---|---|---|
| *Asobara sp.* | AA/A | 0.41 | 0.001 |
| | AL/A | 0.71 | 0.434 |
| | AL/AA | 1.73 | 0.080 |
| | AG/A | 0.70 | 0.082 |
| | AG/AA | 1.70 | 0.082 |
| | AG/AL | 0.99 | 1.000 |
| *Ganaspis sp.* | GG/G | 0.05 | < 0.0001 |
| | AG/G | 0.10 | 0.0002 |
| | AG/GG | 2.02 | 0.183 |
| | LG/G | 0.37 | 0.301 |
| | LG/GG | 7.93 | < 0.0001 |
| | LG/AG | 3.94 | 0.010 |
| *Leptopilina sp.* | LL/L | 0.18 | 0.656 |
| | AL/L | 1.35 | 0.993 |
| | AL/LL | 7.51 | 0.231 |
| | LG/L | 0.51 | 0.931 |
| | LG/LL | 2.81 | 0.768 |
| | LG/AL | 0.38 | 0.124 |

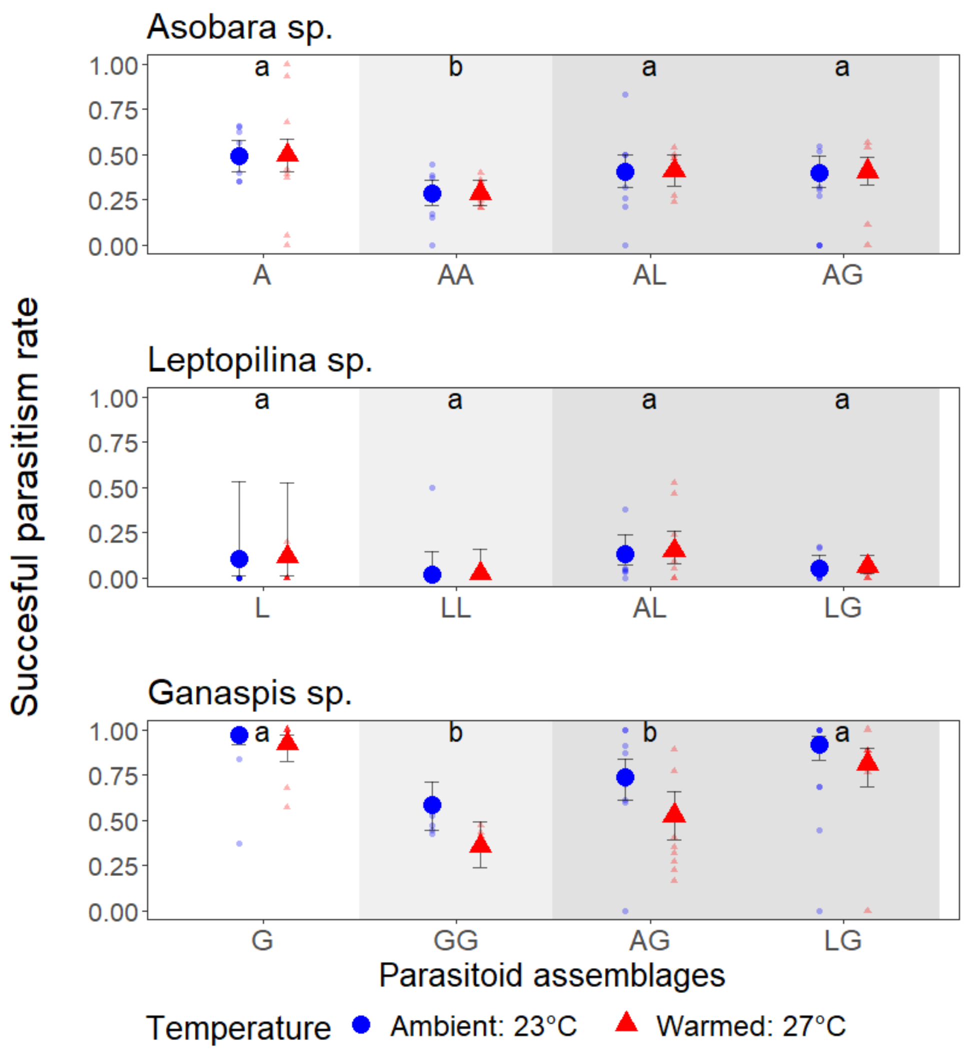


**Figure S5.** Probability of successful parasitism rate varied across parasitoid assemblage and temperature depending on the species identity (N = 144 vials). Different small letters denote significant differences between parasitoid assemblages within each parasitoid species. White panel: single parasitoid, light grey panel: two parasitoids conspecific, darker grey panel; two parasitoids heterospecific. Parasitoid abbreviations: A: *Asobara sp*., L: *Leptopilina sp*., and G: *Ganaspis sp*.. Big dots represent the estimated means (±95% CIs), and small dots represent raw data. Contrasts between parasitoid assemblages are presented in Table S2.

*Supplement Material S7: Effects of warming and parasitoid assemblage on super- and multiparasitism rate*

**Table S11.** Effects of temperature, parasitoid assemblage, and their interactions on the super- and multiparasitism rate. Effects are shown by the summary of Likelihood-ratio chi-square tests with the effects of temperature (2 levels) and parasitoid assemblage (5 levels). Degrees of freedom (Df) are given for each factor and the residuals.

| **Effects** | **$\chi 2$** | **Df** | **p-value** |
|---|---|---|---|
| Temperature | 4.49 | 1 | 0.034 |
| Parasitoid assemblage | 572.40 | 4 | < 0.0001 |
| Temperature: assemblage | 36.04 | 4 | < 0.0001 |
| | | 977 | |

*Supplement Material S8: Effects of warming and parasitoid assemblage on encapsulation frequency*

52.4% of the parasitized larvae (n = 868) had evidence of melanization (traces, melanized egg, and/or melanized larvae), signaling a host immune response. The frequency of encapsulated parasitoids was significantly affected by parasitoid assemblages ($\chi2_{(4)} = 23.89$, $P < 0.0001$), and the interaction between temperature and parasitoid assemblages ($\chi2_{(4)} = 11.42$, $P = 0.0223$), but only because of the difference between *Asobara sp*. and *Ganaspis sp*., not because of parasitoid treatments (single, conspecifics, and heterospecifics; Figure S6). *Asobara sp*. escaped encapsulation due to its fast development time (Figure S3). Indeed, when *Asobara sp*. parasitized larvae, most traces of melanization were observed on the empty eggshell, with the parasitoid larva still alive. Moreover, observations through dissections did not inform us of the outcome of the interactions. Indeed, when *Ganaspis sp*. parasitized larvae, some eggs were only partially encapsulated, and parasitoid larvae could still hatch from these.

**Table S12.** Effects of temperature, parasitoid assemblage, and their interactions on the encapsulation rate. Effects are shown by the summary of Likelihood-ratio chi-square tests with the effects of temperature (2 levels) and parasitoid assemblage (5 levels). Degrees of freedom (Df) are given for each factor and the residuals.

| **Effects** | **χ2** | **Df** | **p-value** |
|---|---|---|---|
| Temperature | 0.002 | 1 | 0.963 |
| Parasitoid assemblage | 23.89 | 4 | < 0.0001 |
| Temperature: assemblage | 11.42 | 4 | 0.022 |
| | | 977 | |

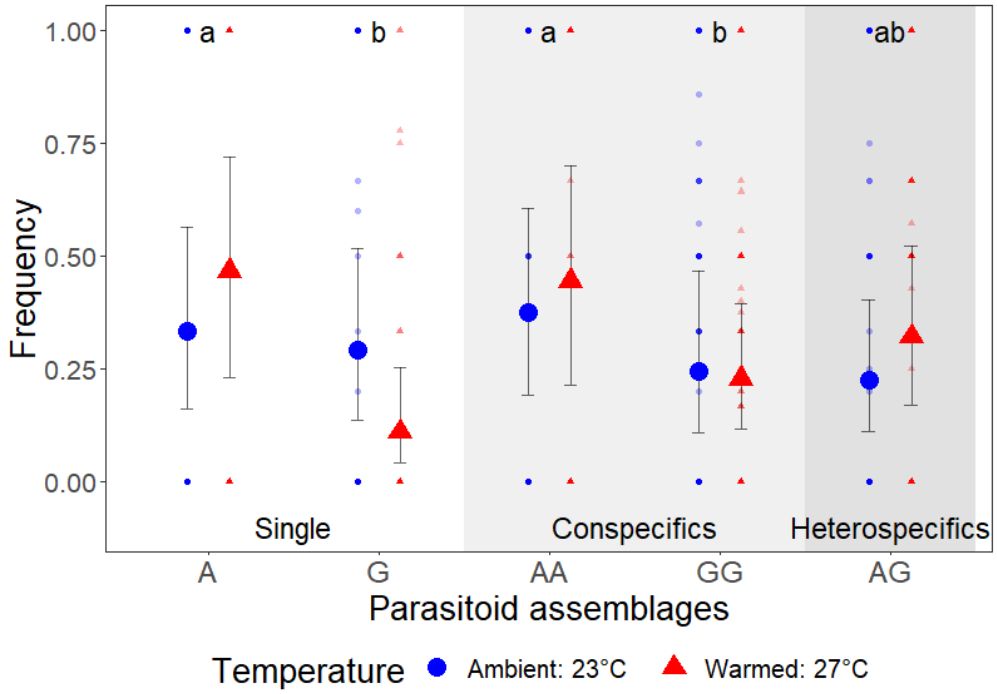


**Figure S6.** Frequency of encapsulated parasitoids out of the total parasitoids per host (N = 1,000 hosts) only changed between parasitoid species. Within each plot, different small letters denote significant differences between parasitoid assemblages. White panel: single parasitoid, light grey panel: two parasitoids conspecific, darker grey panel; two parasitoids heterospecific. Parasitoid abbreviations: A: *Asobara sp*., and G: *Ganaspis sp*. Big dots represent the estimated means (±95% CIs), and small dots represent raw data.
